# Exploring the Antifibrotic Potential of the heparan sulfate mimetic OTR4120: Insights from Preclinical Models

**DOI:** 10.1101/2024.10.17.618798

**Authors:** Manuela Marega, Najet Mejdoubi-Charef, David Wiegard, Muzamil Majid Khan, Marek Bartkuhn, Laura Gambs, Tara Procida-Kowalski, Jochem Wilhelm, Esmeralda Vasquez-Pacheco, Afshin Noori, Ying Dong, Yi Zheng, Xuran Chu, Arun Lingampally, Joanna Zukowska, Said Charef, Franck Chiappini, Agnes Choppin, Dulce Papy Garcia, Rainer Pepperkok, Thomas Muley, Hauke Winter, Clemens Ruppert, Werner Seeger, Andreas Gunther, Denis Barritault, Chengshui Chen, Cho-Ming Chao, Stefano Rivetti, Saverio Bellusci

## Abstract

**Rationale:** Idiopathic pulmonary fibrosis (IPF) is a debilitating lung disease characterized by excessive deposition of extracellular matrix (ECM), resulting in lung function impairment. Heparan sulfate mimetics (HSm) have been suggested to have potential antifibrotic effects by regulating ECM. This study aims to investigate the impact of a specific HSm named OTR4120 on fibrotic processes in *ex vivo*, *in vitro* and *in vivo* models.

**Methods:** Human Precision Cut Lung Slices (hPCLS) treated with a fibrotic cocktail alone or with OTR4120 were evaluated using second-harmonic imaging microscopy (SHIM) to assess collagen deposition. Human embryonic fibroblast WI-38 cell line and primary fibroblasts obtained from human donors were differentiated into myofibroblasts (MYF) using TGF-β1 and treated with OTR4120 or a control vehicle. Gene expression analysis for MYF markers was performed using quantitative PCR (qPCR). Protein expression of MYF markers was evaluated using immunofluorescence techniques. Bulk-RNA sequencing analysis on WI-38 cells cultured under different experimental conditions was conducted. Finally, the therapeutic effects of OTR4120 on a bleomycin-induced fibrosis mouse model were investigated.

**Results:** SHIM analysis on OTR4120-treated hPCLS showed a decrease in collagen deposition. OTR4120 treatment of primary fibroblasts and WI-38 cells exposed to TGF-β1 significantly reduced the expression of MYF markers. Bulk-RNA sequencing analysis on OTR4120-treated WI-38 cells showed significant impacts on fibrosis-related processes. Therapeutic application of OTR4120 *in vivo* to bleomycin-induced fibrosis mice resulted in enhanced fibrosis resolution.

**Conclusion:** OTR4120 has potential therapeutic benefits as an antifibrotic agent in the context of lung fibrosis. Further investigations are necessary to understand the precise mechanism through which OTR4120 exerts its antifibrotic effects.

## Introduction

Pulmonary fibrosis is a progressive and deadly disease characterized by excessive deposition of extracellular matrix components, leading to irreversible scarring of lung tissue, and compromised respiratory function [1]. The etiology of pulmonary fibrosis is multifaceted, encompassing idiopathic forms as well as those associated with environmental exposures, systemic diseases, and infections such as SARS-CoV-2 [1]. Despite advances in understanding the pathophysiological mechanisms underlying this condition, therapeutic options are limited, and the prognosis for affected individuals is poor, with a high mortality rate within five years of diagnosis [1]. Pulmonary fibrosis imposes a substantial burden on affected individuals, impacting their quality of life, functional capacity, and overall survival. As lung tissue progressively scars, patients experience dyspnea, cough, fatigue, and reduced exercise tolerance, severely limiting daily activities and often causing the need of supplemental oxygen therapy. The therapy is limited, and currently only two drugs, pirfenidone and nintedanib, are approved for IPF [2, 3]. Unfortunately, both pharmacological agents only slow down the progression of the disease but fail to stop it.

In the search for more effective treatments, the antifibrotic agent OTR4120 has emerged as a potential candidate. OTR4120 belongs to a class of ReGeneraTing Agents (RGTA®), designed to improve the natural regenerative processes [4]. RGTA® are polysaccharides specifically engineered to mimic heparan sulfate, a key component of the extracellular matrix (ECM) that plays a crucial role in maintaining tissue homeostasis [5]. Heparan sulfate proteoglycans (HSPG) are integral to the ECM, where they regulate a wide range of biological processes by binding to a diverse panel of proteins, including growth factors, cytokines, and enzymes [6]. The degradation of HSPG following tissue injury disrupts this delicate equilibrium and may impair the wound healing process. The application of the RGTA® in the clinical setting covers different area, such as wound healing and treatment of corneal lesions [7], as well as stroke consequences and ischemic lesion [8].

This paper aims to explore the potential antifibrotic effects of OTR4120 in lung fibrosis. Due to its successful action in the wound healing, with a significant reduction of the scars [5, 9], we hypothesized that OTR4120, with its potential ECM remodelling features, could exert a therapeutic action in the context of lung fibrosis.

The unique properties of RGTA®, such as resistance to degradation and the ability to protect ECM structural and signalling proteins, like HSPG, offer a promising avenue for restoring both structural and biochemical functions to the ECM, thereby helping the processes of tissue repair and regeneration.

In this study, we investigated the effect of OTR4120 treatment *ex vivo* on human Precision Cut Lung Slices (hPCLS) from donor lungs, *in vitro*, on human primary fibroblasts isolated from donor lungs and on a human embryonic lung fibroblast cell line called WI-38 [10] as well as *in vivo* in a bleomycin-induced mouse model of pulmonary fibrosis [11, 12].

OTR4120 has potential therapeutic benefits as an antifibrotic agent in the context of lung fibrosis but further investigations are necessary to understand the precise mechanism through which OTR4120 exerts its antifibrotic effects.

## Materials and methods

### hPCLS *ex vivo* culture and label-free SHG imaging

Tumor-free lung tissue was obtained from Thoraxklinik-Heidelberg, Germany, with patient consent and ethical approval (S-270/2001, BIAC 2021-003). Precision-cut lung slices (PCLS) were prepared from the tissue samples and treated with a fibrotic cocktail to induce fibrosis. Subsequently, the PCLS were treated with OTR4120 and analyzed using Second-Harmonic imaging microscopy (SHIM) following established protocols [13]. Image analysis involved calculating the total fibrillar collagen content based on Second-Harmonic Generation (SHG) signal intensity, with results normalized to control samples from the same donor. Details in the supplementary information file.

### Cell Lines and cell culture

Human primary lung fibroblasts (HP-LFs) were obtained with patient consent from the European IPF registry, and the WI-38 cell line was sourced from ATCC. Both cell types were cultured under standard conditions and subjected to treatments, including TGF-β1, OTR4120, and SB431542. Detailed cell culture conditions and treatment protocols are provided in the supplementary material.

### Immunofluorescence

See supplementary material.

### RNA extraction and qPCR

See supplementary material.

### FACS preparation and alveolosphere assay

See supplementary material.

### Statistical analysis

Statistical analyses and graph assembly were conducted using GraphPad Prism (v6.0) (GraphPad Prism Software). To compare the means of two groups, an unpaired, two-tailed Student’s t-test was used. For comparing the means of three or more groups, a one-way ANOVA with post hoc analysis was done. Assessment for the presence of outliers was performed using ROUT analysis. The figure legends include the number of biological samples (n) for each group and the specific statistical tests used. Type I error was set at 5% (i.e. α= 0.05). Statistical significance was considered if the p-value was less than 0.05.

### RNA sequencing and bioinformatic analysis

RNA sequencing of WI-38 cells was performed, followed by bioinformatic analysis to assess genome-wide gene expression. Libraries were prepared, sequenced, and processed using standard protocols, with differential gene expression analyzed in R. Detailed methods, including library preparation, sequencing, and bioinformatics processing, are available in the supplementary material.

RNA-seq data has been deposited at NCBI’s gene expression omnibus (GEO) the accession is GSE279215.

### Silencing of *LBH*

See supplementary material.

### in vivo model

Lung fibrosis was induced in male Swiss mice using bleomycin delivered intraperitoneally, followed by treatment with OTR4120 or saline delivered intravenously. Mice were monitored for 28 days, with assessments of fibrosis, inflammation, and tissue remodeling. All animal procedures were conducted in accordance with established ethical guidelines, with protocol approval granted by the Animal Care Committee of French University in Paris, France (APAFIS# 19838). Detailed protocols, including the treatment schedule and evaluation criteria, are provided in the supplementary material.

## Results

### OTR4120 reduces fibrillar collagen deposition in ECM in hPCLS treated with a fibrotic cocktail

First, we investigated the effect of OTR4120 on fibrillar collagen deposition in a physiological fibrogenesis model using the *ex vivo* hPCLS culture system [13] (Figure 1a). Excessive fibrillar collagen deposition only occurs when a pan metalloproteinase inhibitor (Ilomastat) is co-administered with TGF-β1 [13]. To replicate the fibrosis process and induce excessive fibrillar collagen deposition, hPCLS were cultured with a fibrotic cocktail (FC) containing TGF-β1 and Ilomastat. The hPCLS remained viable for at least 2 weeks under the established conditions [13]. After two weeks of FC treatment, the deposition of fibrillar collagen in the ECM was quantified using label-free SHG imaging. The analysis revealed a significant increase in fibrillar collagen deposition compared to control samples, confirming the pro-fibrotic effect of the FC (Figure 1b, c). Next, we examined the impact of OTR4120 on fibrillar collagen deposition in the hPCLS. The hPCLS were treated with OTR4120 in the presence of the FC for the same duration (Figure 1d). Remarkably, the deposition triggered by the FC was significantly reduced when OTR4120 was present (Figure 1e, f). Similarly, qPCR results obtained from these hPCLS after 48h (Figure S1a) confirm the significant decrease in *ACTA2* and *COL1A1* expression upon OTR4120 treatment (Figure S1b and c, respectively). These data demonstrate the potential anti-fibrotic action of OTR4120 in this *ex vivo* fibrogenesis model.

**Figure 1:**
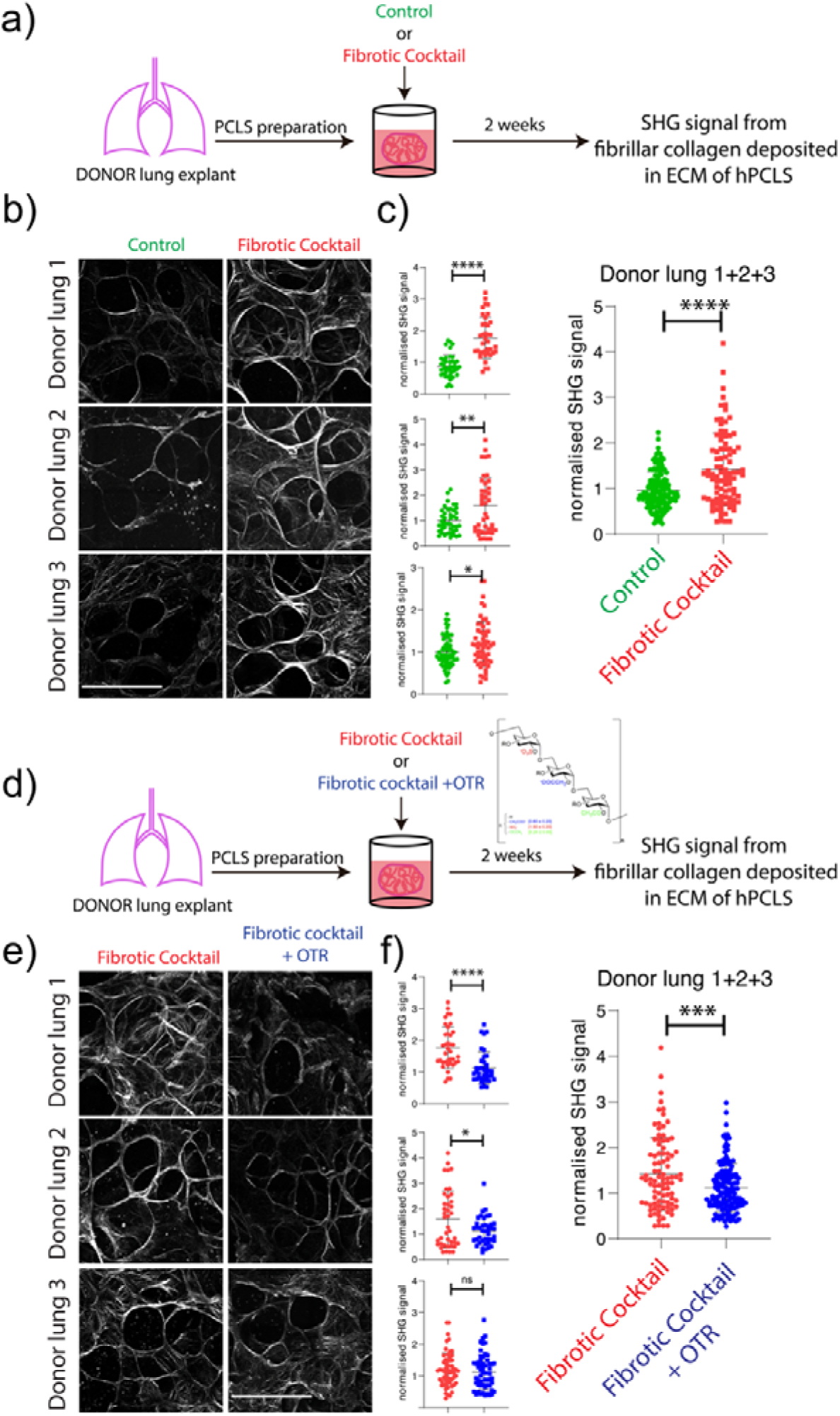
OTR4120 treatment of PCLS from human DONOR lungs grown in vitro in presence of a Fibrotic Cocktail leads to reduced Collagen deposition. **(a)** Schematic representation of hPCLS treatment with Fibrotic Cocktail (FC). **(b)** Representative images of three biological replicates: left panel shows control, right panel shows FC treatment, after 2 weeks, with corresponding quantification. **(c)** Average measurements from the quantification in (b). **(d)** Schematic representation of hPCLS treatment with FC and FC/OTR4120 (simplified as OTR in the figure). **(e)** Representative images of three biological replicates: left panel shows control, right panel shows FC treatment, after 2 weeks, with corresponding quantification. **(f)** Average measurements from the quantification in (e). Scale bar b-e: 50 μm. P values * P < 0.05; ** P < 0.01; *** P < 0.001; **** P < 0.0001.

### OTR4120 treatment decreases the expression of fibrotic markers in primary human lung fibroblasts

To specifically investigate the cell type targeted by OTR4120 antifibrotic effect, we used primary human lung fibroblasts (HP-LFs) derived from donors from different ages and gender (Figure 2a). It has been previously documented that upon treatment with TGF-β1, HP-LFs undergo differentiation into myofibroblasts (MYF), which are responsible for excessive ECM deposition during lung fibrosis [14]. To assess the effect of OTR4120 on myofibroblasts, cells were first treated with TGF-β1 for 96h to induce myofibroblast differentiation and subsequently treated with OTR4120 or vehicle for an additional 96h (Figure 2b). Our results indicated that, compared to the vehicle, OTR4120 treatment resulted in a decrease in *ACTA2* and *MHY11* expression (Figure 2c). These findings strongly indicate that OTR4120 directly acts on myofibroblasts, prompting their dedifferentiation. The specific identity of this fibroblast subtype remains to be determined and requires further investigation.

**Figure 2:**
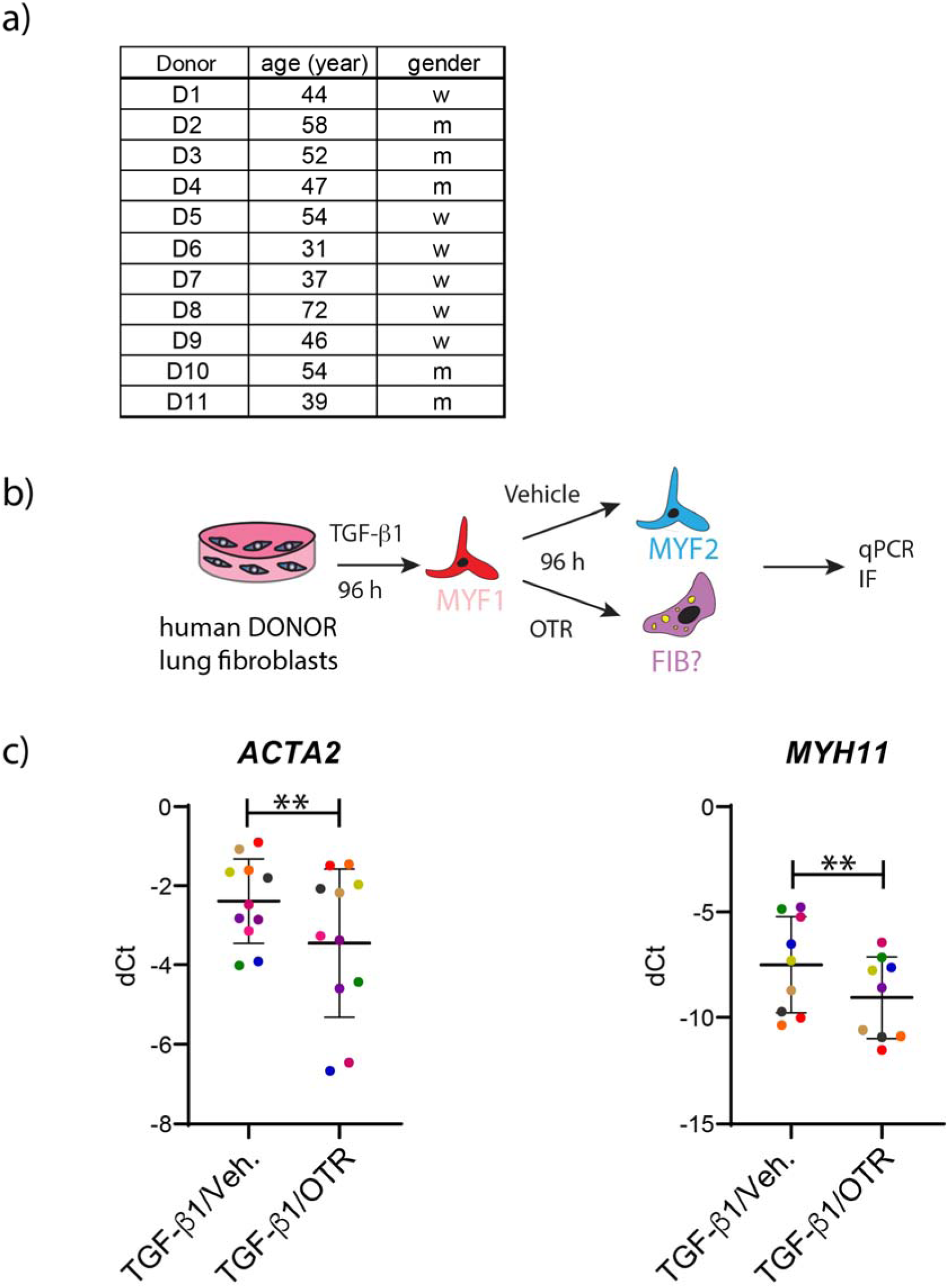
OTR4120 treatment of primary fibroblasts of human DONOR lungs leads to reduced fibrotic markers expression. **(a)** Schematic representation of the human primary lung fibroblast (HP-LFs) experimental set up. **(b)** qPCR for MYF markers: *ACTA2* (left), *MYH11* (right). (c) List of the healthy donor for HP-LFs (age and gender). P values * P < 0.05; ** P < 0.01; *** P < 0.001; **** P < 0.0001.

### OTR4120 decreases the expression of fibrotic markers in TGF-**β**1-treated WI-38 cells and promotes alveolo-sphere formation

To dive deeper into the potential antifibrotic effect of OTR4120 on myofibroblasts, the WI-38 cell line was used as an *in vitro* model to characterize the impact of the drug [10]. WI-38 cells were treated similarly to the primary cells (Figure 3a).

**Figure 3:**
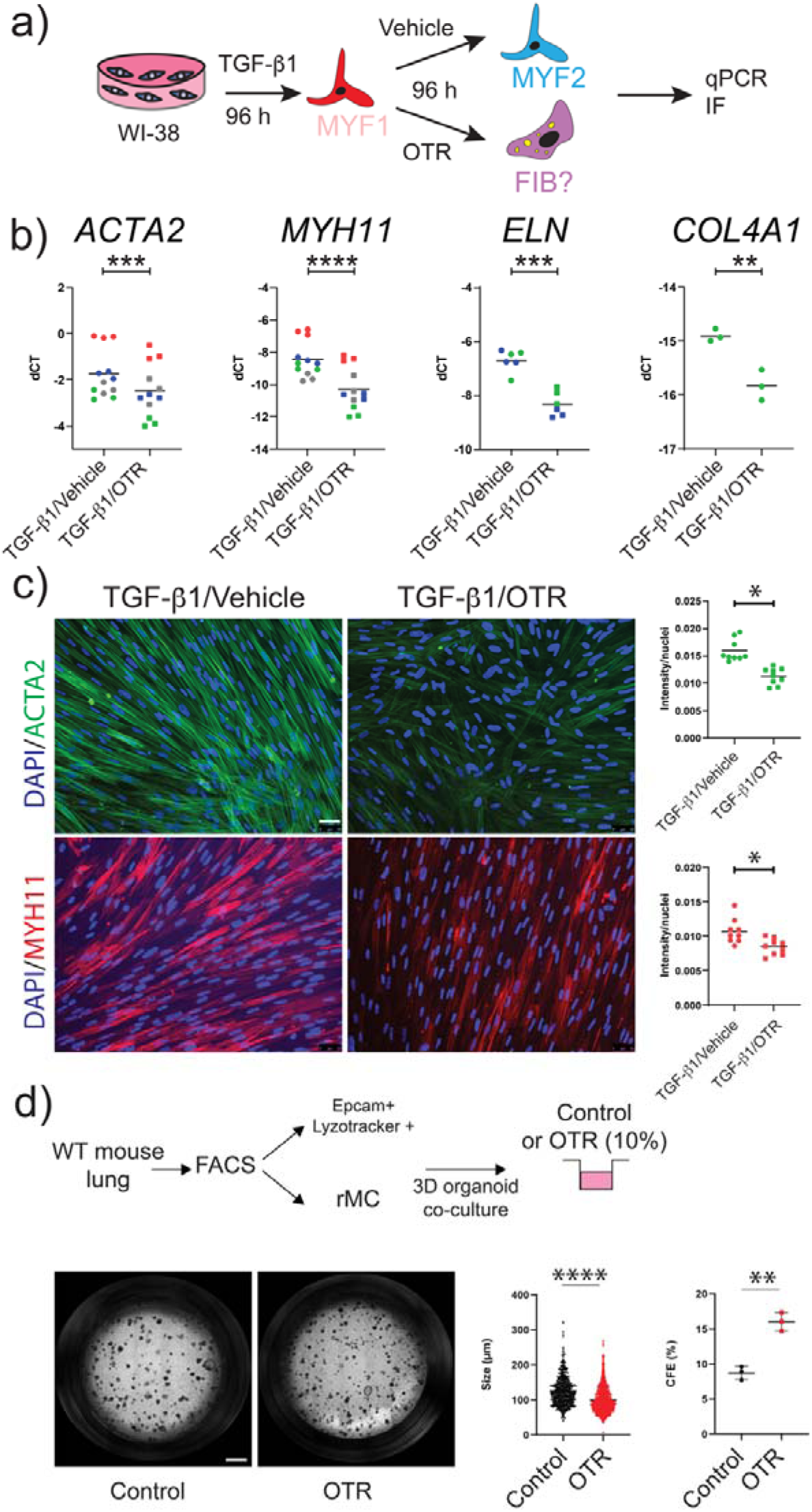
OTR4120 treatment of primary fibroblasts of WI-38 cells leads to reduced fibrotic markers expression. **(a)** Schematic representation of the WI-38 experimental set up. **(b)** Gene expression analysis for the MYF markers: *ACTA2*, *MYH11*, *ELN* and *COL4A1*. **(c)** Immunofluorescence for ACTA2 and MYH11 (ctrl and OTR4120) (left), quantification of the signal (right). Nuclei are stained with DAPI. **(d)** Alveolosphere assay: schematic of the experimental set up, representative images for control and OTR4120-treated samples (left), measurement of the size and CFE (%) of the spheres (right). *P* values * *P* < 0.05; ** *P* < 0.01; *** *P* < 0.001; **** *P* < 0.0001. Scale bar c and d: 50 μm and 800 μm.

WI-38 cells were first treated with TGF-β1 for 96h to induce their differentiation towards the MYF lineage. Analysis by qPCR and IF at this time points show increased in MYF markers compared to the vehicle-treated cells (Fig S2b). Following TGF-β1 treatment, MYF were cultured for an additional 96h with either vehicle or OTR4120.

Analysis of markers for myofibroblast (MYF) differentiation induced by the TGF-β1 treatment, such as *ACTA2*, *MYH11*, *ELN*, and *COL4A1*, showed a significant decrease in gene expression following OTR4120 treatment compared to the vehicle group (Figure 3b). Consistent with this, protein levels of ACTA2 and MYH11 were also reduced in IF, confirming the antifibrotic effect of OTR4120 in the cells (Figure 3c). Interestingly, in the alveolo-sphere assay, when WI-38 and alveolar type 2 (AT2) cells were mixed, OTR4120 treatment impacted positively the initiation of organoid formation with a significant increase in CFE (Figure 3d), suggesting that OTR4120 could be potentially beneficial *in vivo* to enhance the repair process.

### OTR4120 action is via TGF-β1 pathway, but not limited to it: preventive and synergistic approach

Based on the observed strong inhibition of MYF differentiation markers, we decided to investigate the specific effect of OTR4120 on the TGF-β1 pathway. To do so, we conducted a preventive approach experiment using WI-38 cells, where OTR4120 was administered concomitantly with TGF-β1 (Figure 4a). The cells were treated with vehicle, TGF-β1 alone or TGF-β1 in combination with OTR4120. Remarkably, the presence of OTR4120 was sufficient to prevent MYF differentiation. *ACTA2* and *MYH11* were significantly lower in the combination group compared to cells treated with TGF-β1 only (Figure 4b), evidence confirmed by the IF analysis for ACTA2 (Figure 4c). To gain further insights into the mechanism of action and explore alternative pathways involved in fibrosis, we compared the effects of SB431542, a TGF-β1 receptor inhibitor, and OTR4120 on WI-38 [15, 16]. Cells were subjected to pretreatment with TGF-β1 and then exposed to either the vehicle, SB431542 alone, OTR4120 alone, or a combination of the two drugs (Figure 4d). Both SB431542 and OTR4120 individually were able to reverse the MYF phenotype, as indicated by the significant decrease in gene expression of *ACTA2* and *MYH11*. Intriguingly, the combination treatment further impacts these markers, with a trend towards a decrease for *ACTA2* and a significant decrease for *MYH11*, suggesting that OTR4120 acts, not only through the TGF-β1 pathway, but also involves additional pathway(s) (Figure 4e). IF analysis revealed a reduction in ACTA2 expression, which was particularly prominent when both drugs were present (Figure 4f), similarly to what we observed at the transcriptional level. Taking these data together, OTR4120 acts on TGF-β1 pathway, but its mechanism of action is not limited to this specific pathway. It acts synergistically with the TGF-β1 receptor inhibitor SB431542, highlighting its ability to target alternative pathways involved in the fibrotic process.

**Figure 4:**
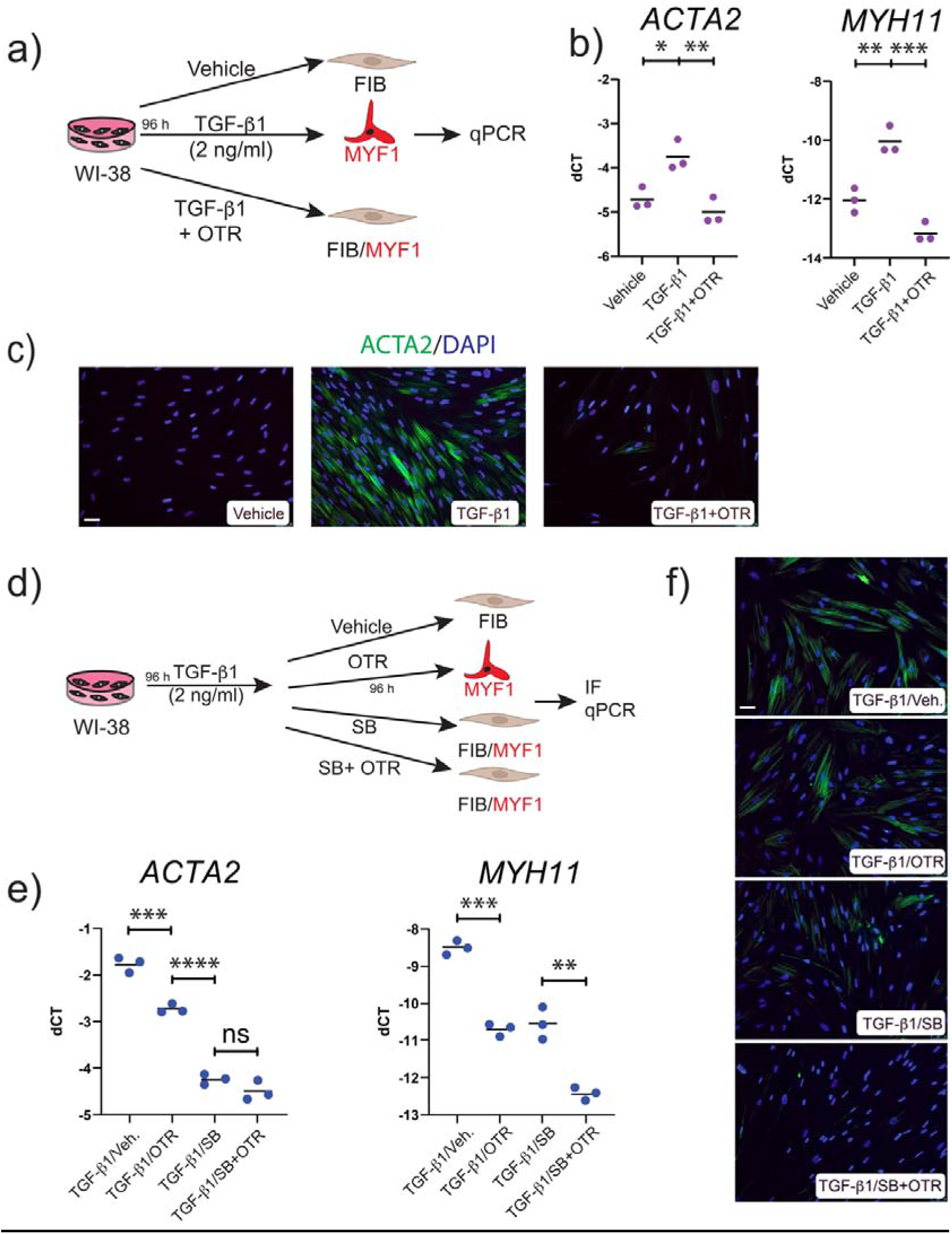
Evidence in WI-38 cells for the efficiency of OTR4120 in a preventive in vitro protocol and for a synergistic antifibrotic effect with TGF-β1 inhibitor. **(a)** Schematic representation of the experimental set up for the preventive approach. **(b)** Gene expression analysis for the MYF markers: *ACTA2* (left) and *MYH11* (right). **(c)** Immunofluorescence for ACTA2 and MYH11 in the three different conditions (vehicle, TGF-β1-treated, and OTR4120). Nuclei are stained with DAPI. **(d)** Schematic representation of the experimental set up for the synergistic approach. **(e)** gene expression analysis for the MYF markers: *ACTA2* (left) and *MYH11* (right). **(f)** Immunofluorescence for ACTA2 and MYH11 for the four different conditions (vehicle, SB431542-treated only, and OTR4120-treated only, combinatorial treatment SB431542/OTR4120. *P* values * *P* < 0.05; ** *P* < 0.01; *** *P* < 0.001; **** *P* < 0.0001. Scale bar c, f: 50 μm.

### Bioinformatic analysis reveals alternative pathway for OTR4120 mode of action

We performed a bulk RNA-Seq analysis to further investigate OTR4120 effects in the fibrotic process and we used our previously described *in vitro* WI-38 cell model (Figure 5a). Bulk RNA-Seq for the OTR4120-treated group (TGF-β1/OTR4120) versus vehicle group (TGF-β/Vehicle) were carried out to assess the overall impact of OTR4120 treatment. Principal Component Analysis (PCA) demonstrated that OTR4120-treated cells segregated distinctly from the vehicle control group, with the first principal component (PC1) accounting for 71% of the variation (Figure 5b). Next, we generated a heatmap to visualize the differentially expressed genes (DEGs) (Figure 5c). In the heatmap, downregulated genes included important MYF related genes such as *MYH11*, *SERPINE1*, *GREM1*, and *GREM2* as well as upregulated genes including *PTGDS* (*prostaglandin D2 synthase*), *MATN2* (*Matrilin-2*) and *SPOCK1* (*Testican-1*) (Figure 5c) providing additional support for the antifibrotic effects of OTR4120 and suggesting a complex mechanism of action. Gene Set Enrichment Analysis (GSEA) was performed to assess the enrichment of TGF-β-induced genes between the OTR4120 condition and the control. Gene Set Enrichment Analysis (GSEA) was performed to assess the enrichment of TGF-β-induced genes between the OTR4120 condition and the control. The TGF-β1 signature, from our previous publication [10] shows a negative trend, with the Normalized Enrichment Score (NES) being negative, indicating that TGF-β-induced genes are underrepresented in OTR4120 samples versus control. This is statistically significant, with both a nominal p-value and adjusted p-value (FDR q-value) of 0.007578. The ranked list metric further highlights the concentration of TGF-β-induced genes in the lower ranks of the dataset, reinforcing the observed negative enrichment and suggesting suppression of TGF-β-induced pathways in OTR4120, likely reflecting a diminished response to TGF-β signaling compared to control. The downregulation of the TGF-β1 signature confirms the impact of OTR4120 on this pathway (figure 5d). Additionally, the bar plot illustrates the top Reactome pathways identified through GSEA. Both extracellular matrix (ECM) organization and immune-related pathways (such as interleukin-4 and interleukin-13 signaling) show negative NES, indicating downregulation in OTR4120 (Figure 5e). This suggests a reduction in ECM-related gene expression, potentially reflecting decreased ECM deposition or remodeling, and suppressed immune activity, particularly in pathways involved in tissue repair and immune regulation. The simultaneous downregulation of both ECM-related and immune-related pathways suggests that the OTR4120 condition involves a globally reduced inflammatory and tissue remodeling profile. These findings could be critical for understanding the molecular mechanisms at play in OTR4120, particularly in the context of fibrosis and immune regulation.

**Figure 5.**
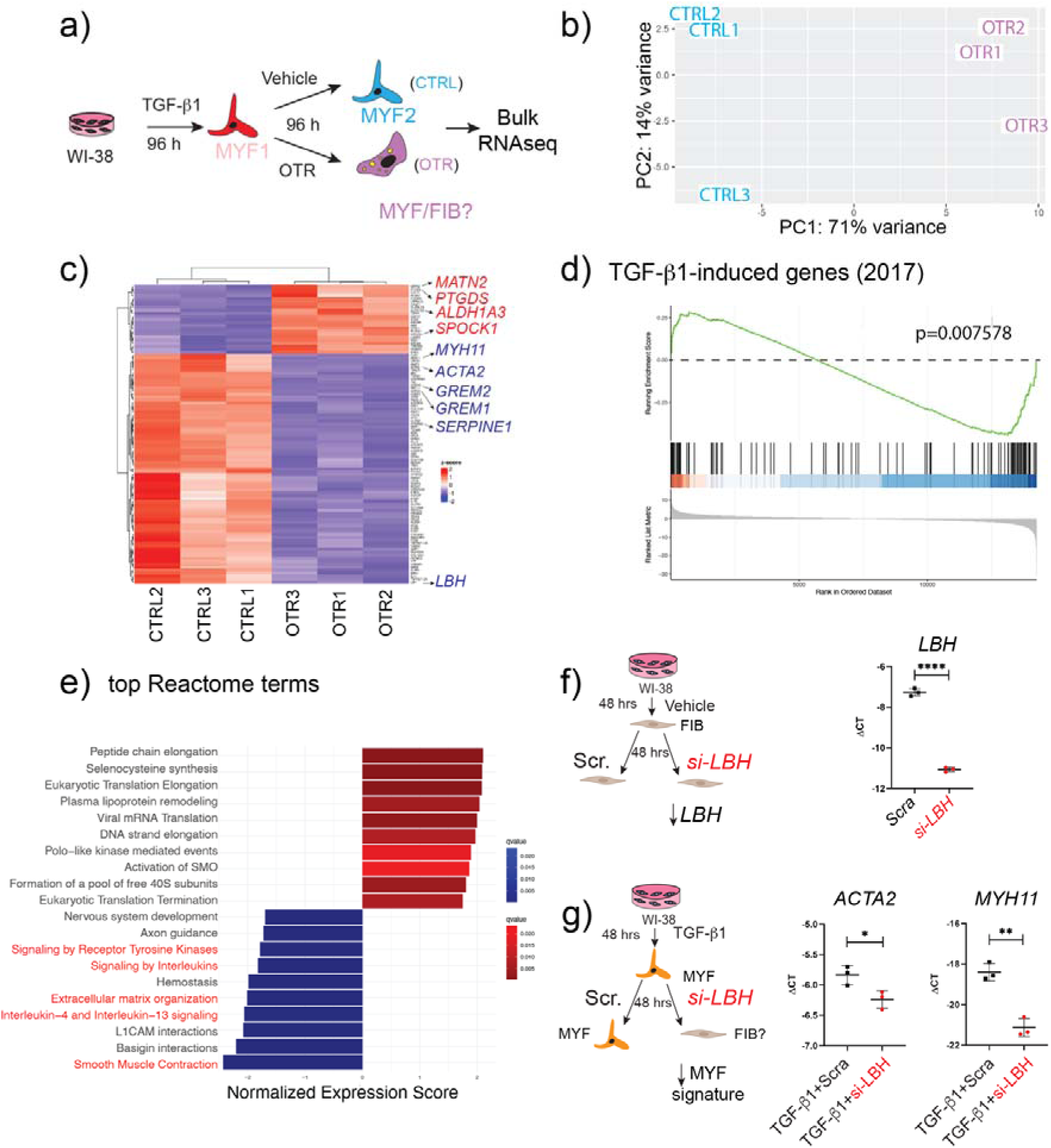
Analysis of transcriptomic changes upon OTR4120 treatment support the inhibition of TGF-β1 signalling. **(a)** Schematic representation of the experimental set up (treatment). **(b)** Principal Component Analysis. **(c)** Heatmap. **(d)** GSEA analysis of TGF-β-induced genes in OTR vs. control. **(e)** GSEA of top Reactome pathways in OTR vs. control. **(f)** Schematic representation of the silencing for *LBH*. Scrambled RNA is used as control. Corresponding validation by qPCR of the decrease in *LBH* expression. **(g)** Schematic representation of the silencing for *LBH* in the context of TGF-β1 treatment. Corresponding impact on *ACTA2* and *MYH11* expression. *P* values * *P* < 0.05; ** *P* < 0.01; *** *P* < 0.001; **** *P* < 0.0001.

These findings underline the potential of OTR4120 in promoting tissue regeneration, attenuating the ECM deposition and promoting healthy remodeling, supported by an action on the immune compartment through reduction of the inflammation. Overall, our bulk RNA-Seq analysis provides strong evidence of the antifibrotic and regenerative effects of OTR4120. It confirms the modulation of the expression of the typical MYF markers, but also sheds light on the fact that OTR4120 can modulate the MYF signature to promote tissue regeneration and repair.

Interestingly, *LBH* (*Limb Bud Heart*) is one of the top downregulated genes in the OTR4120 treatment (Figure 5c). *LBH* encodes a transcription co-factor in the Wnt/β-catenin pathway, that is upregulated by TGF-β1 and involved in the activation of the myofibroblast in fibrotic diseases [17, 18]. Heparan sulfate glycans are known to modulate the Wnt pathway and the likely alteration of Heparan Sulfate components could impact directly or indirectly LBH protein function.

The silencing of *LBH* in untreated WI-38 was validated (Figure 5f). In TGF-β1 induced MYFs, the knocking down of *LBH* decreased the expression of the MYF markers *ACTA2* and *MHY11* (Figure 5g). This result suggests that the decrease in *LBH* expression upon OTR4120 treatment, likely due to the general inhibition of TGF-β1 signaling, is functionally responsible for the observed antifibrotic effect.

### OTR4120 accelerates fibrosis resolution following bleomycin administration in mice

Next, we carried out a preclinical model to study the effect of OTR4120 *in vivo*. The bleomycin-induced lung fibrosis model in mice is the gold standard model to evaluate the overall effect antifibrotic effect of OTR4120. We investigated the effects of OTR4120 treatment on bleomycin-induced lung fibrosis in mice, using control groups (Saline Solution and OTR4120 only), and experimental groups, where the mice received bleomycin intraperitoneally, followed by either saline solution, or OTR4120 administration (I.V.) (Figure 6a). The mice were sacrificed at day 28 and the lungs analysed. The lung tissue sections from the control groups (Saline solution or OTR4120 only) showed normal alveolar spaces, normal thickening of the alveolar septa and high elasticity (Figure 6b, upper left and right panel). Lung sections from mice in the bleomycin-induced fibrosis group showed marked histopathological abnormalities, with alveolo-interstitial inflammation, at day 28 (Figure 6b, bottom left panel). Lung sections from mice treated with OTR4120 after instillation of bleomycin (Bleo) showed moderate reduction of inflammatory cell infiltration, with normal alveolar structure and a few macrophages, lymphocytes, and plasma cells (Figure 6c, bottom right panel). Furthermore, lung index values (quantitative lung index, QLI) [19], which show that lung damage were significantly lower in the (Bleo/OTR4120)-treated mice compared to the Bleo-injured mice (Figure 6c, left panel). Alveoli Area scores which reflect the presence of functional respiratory tissue, were significantly higher in the (Bleo/OTR4120)-treated mice compared to the Bleo mice. However, the OTR4120-treated group still displayed a reduction compared to the control groups indicating the presence of residual fibrosis (Figure 6c, right panel).

**Figure 6.**
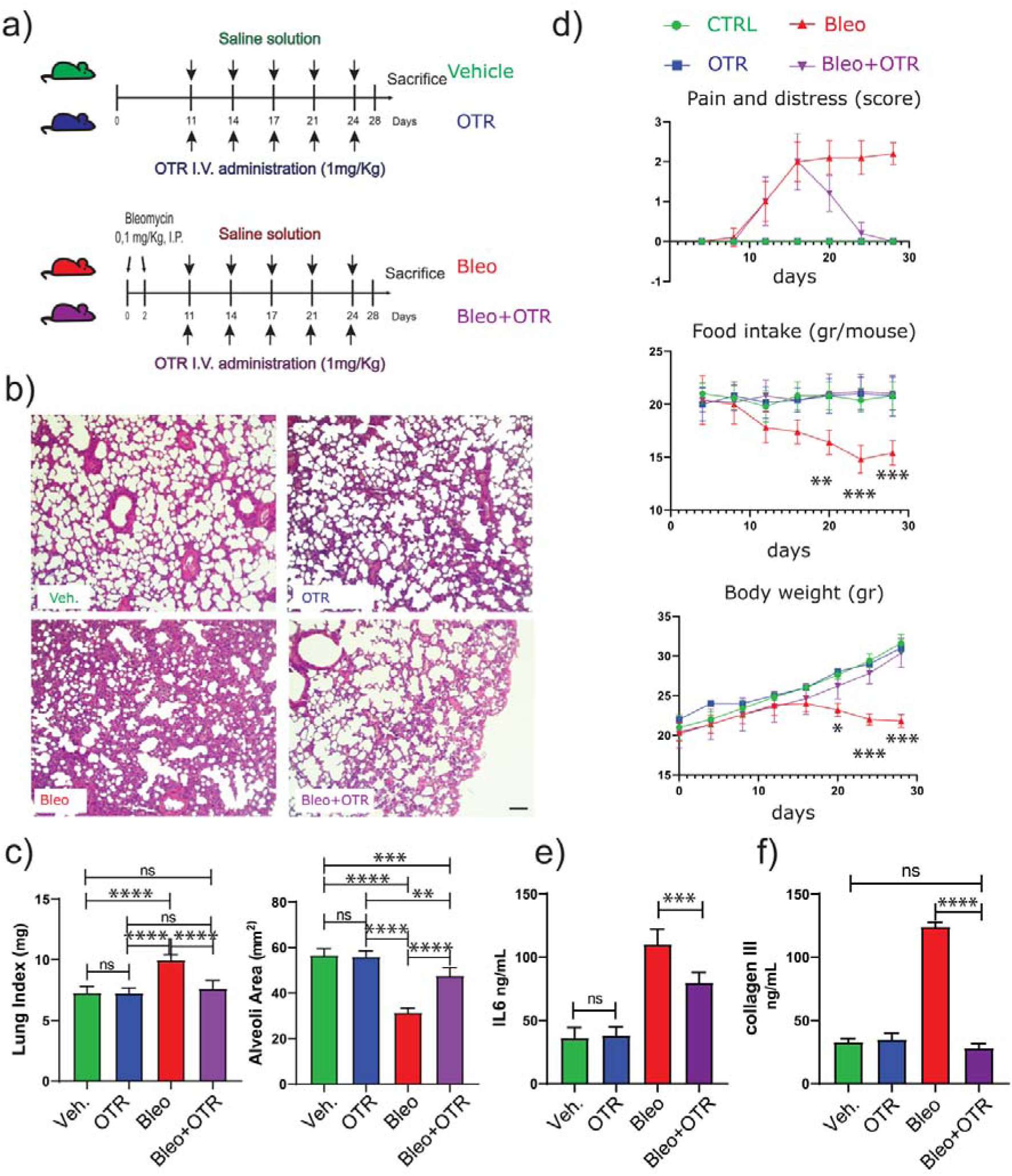
OTR4120 treatment attenuates bleomycin-induced fibrosis in vivo. **(a)** Schematic illustration of the experimental design for the animal study: the groups received only saline solution or only OTR4120 (upper panel) and Bleo/saline solution or Bleo/OTR4120 (lower panel). **(b)** Representative photomicrographs showing haematoxylin and eosin (H&E) stained lung sections for all the four groups: saline only (left panel up), OTR4120 only (right panel up), Bleo/saline solution-treated (left panel down), and Bleo/OTR4120-treated (right panel down). **(c)** Lung Index (left) and Alveoli Area (right) measurements. **(d)** Scores for Pain and distress, Food Intake and Body weight for all the experimental groups. **(e)** Il6 (left) and collagen II (right) measurements in all the experimental groups. Scale bar b: 40 μm.

After 17 days of treatment (from day 11 to 24), the mice treated with OTR4120 showed a reversal of clinical signs associated with bleomycin administration which included pain and distress, food intake and body weight reductions (Figure 6d). Strikingly, the mice that received OTR4120 treatment after induction of fibrosis were comparable with the healthy mice that received saline solution only and OTR4120 alone, indicating that the drug is well-tolerated by the animals, with minor or no side effects. We also measured the expression of the pro-inflammatory cytokine Il-6 in the bronchoalveolar lavage of the mice. Il-6 protein level is reduced in the OTR4120-treated Bleo-injured mice, strongly advocating for an additional effect on the inflammatory process driven by OTR4120 (Figure 6e). The presence of Col3a1 was also reduced in the lung homogenates (Figure 6f).

## Discussion

RGTA® are a class of medicinal substances, which action enhances both the speed and quality of tissue healing, and, in some cases, leads to pronounced tissue regeneration. They interact and protect cellular signalling proteins—such as growth factors, cytokines, interleukins, colony-stimulating factors, chemokines, and neurotrophic factors—from proteolytic degradation to keep tissue homeostasis. These proteins are crucial for cellular communication and are naturally stored in the extracellular matrix, where they specifically interact with heparan sulfate (HS) [20]. RGTA® are in preclinical and clinical stages for the treatment of different kind of organ injuries, that include the brain, where they are tested in the context of acute ischemic stroke and in the subsequent follow up management of the associated symptoms (clinical phase - NCT04083001 [21]).

Moreover, other disease areas where RGTA® could be effective are diseases associated with metabolic syndromes (Type 2 Diabetes, diabetic ulcers, clinical study NCT01474473 [22]) where OTR4120 is in post-market clinical follow up [23]) and dermatology (recruitment phase for clinical studies for post-surgical scars (clinical study NCT05528328 -Phase 3)), as well as in the treatment of epidermolysis bullosa (clinical study NCT06007235). Additionally, RGTA® are also effective in the repair of corneal lesions [24].

Prior to this work, OTR4120 was tested on a compassionate basis in a limited cohort of 13 COVID-19 patients [25]. All the patients improved significantly from the clinical point of view within one month of treatment onset. CT scan highlighted an improvement of the lesions in the lung as well [25]. This study, although limited due to the absence of a control group, opened the way for the use of OTR4120 as a new therapeutic option for pulmonary diseases that involve tissue remodelling, such as lung fibrosis. Therefore, based on these results, we decided to test the efficacy of the OTR4120 in the context of lung fibrosis using *ex vivo* (hPCLS) and *in vivo* (bleomycin injury in mice), as well as *in vitro* (HP-LF and WI-38 cells) models. The results of our studies provide strong evidence for the antifibrotic properties of OTR4120 in all the approaches. In the *ex vivo* hPCLS, OTR4120 effectively reduced collagen deposition, a characteristic feature of fibrotic tissue remodelling. This finding is extremely significant due to the closely replicated microenvironment of human lung tissue in the hPCLS model [26]. Diving deeper in the validation of this antifibrotic effect, OTR4120 was also evaluated on HL-PFs. OTR4120 treatment reduced fibrotic marker expression in HL-PFs, showing its specificity in targeting MYFs, the cell type that plays a central role in fibrosis development. Similarly, our established *in vitro* WI-38 model for fibroblast differentiation [10] further supports the antifibrotic effects of OTR4120, as it decreased the expression of MYF differentiation markers similarly to the HL-PFs. OTR4120 shows potential as a drug for promoting alveolo-sphere initiation, indicating its potential usefulness in lung repair and regeneration. To further explore its potential, it should be tested in the context of elastase injury-induced alveolar destruction or pneumonectomy. By examining the effects of OTR4120 on *de novo* alveologenesis in these two experimental models, we could gain a clearer understanding of its impact on lung regeneration. The dual action of OTR4120, antifibrotic and promoting repair, represents a novel therapeutic concept for fibrosis treatment that requires further investigation.

The proposed mechanism of action of OTR4120 is shown in figure 7, where it is suggested that OTR4120 repairs the damaged extracellular matrix. This action leads to a reduction of TGF-β1 signalling and increase in the bioavailability of endogenous growth factors. Additionally, OTR4120 may act directly on the cells themselves, although further studies are needed to narrow down its specific mechanism of action. Our bioinformatics results demonstrated a decrease in the overall growth factor pathway score following OTR4120 treatment. On the other hand, we observed a decrease in TGF-β1 OTR4120 treatment, suggesting that the main mechanism of action of OTR4120 is through the inhibition of TGF-β1 signalling. This inhibition may occur by preventing the release of latent TGF-β1 or enhancing the binding to ECM/OTR4120 to TGF-β1. OTR4120 binds and protects from proteolytic degradation many communication peptides and our data could result from sequester, blockage or increase of the quantities of active molecules, as shown on other *in vitro* [27] and *in vivo* [28]. The delay between addition of OTR4120 and measures of the induced effects should be parameters to separate direct from indirect effects.

**Figure 7.**
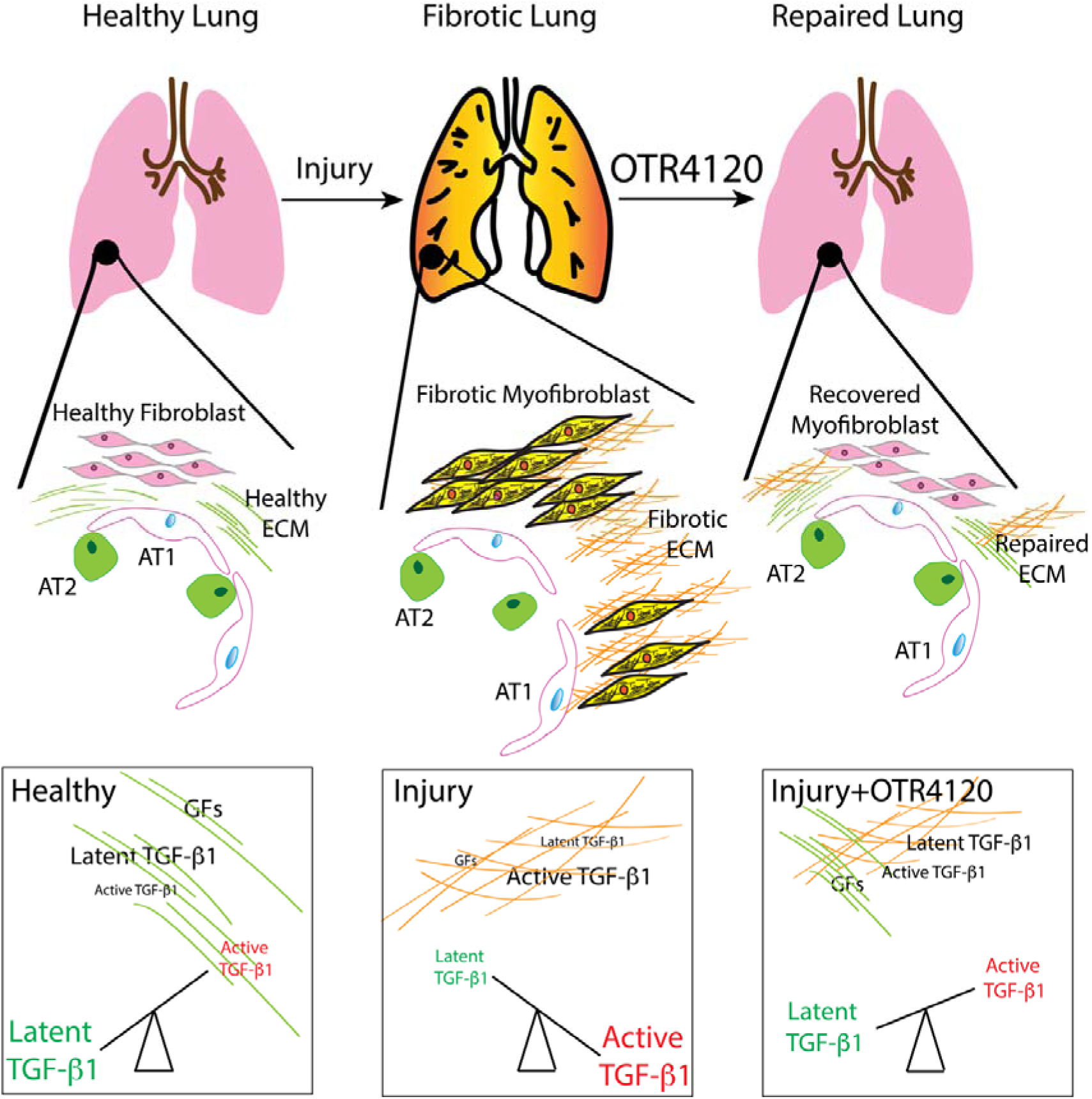
Graphical representation of the action of the OTR4120. Healthy lungs upon injury can develop fibrosis. The fibrotic process is characterized by the differentiation of fibroblast towards activated myofibroblasts leading to increased deposition of dysregulated (fibrotic) extracellular matrix (ECM). OTR4120 treatment reverses this differentiation process, triggering activated myofibroblasts differentiation towards a healthy fibroblast status (called recovered MYF in the figure). The ECM returns to a healthy status compatible with alveolar epithelial repair. The balance between latent TGF-β1 and active TGF-β1 is dependent on the status of the ECM, Healthy or repaired ECM leads to an decrease in the latent active TGF-β1 to TGF-β1 ratio while pathological/injured ECM leads to the corresponding increase in this ratio.

The downregulation of the transcription factor LBH reenforces the hypothesis of the mechanism of action through the TGF-β1 pathway [29]. However, OTR4120 action is not limited to the TGF-β1 pathway as suggested using the pharmacological inhibitor SB431542 for the TGF-β1 signalling pathway. The synergic effect observed on *MYH11* and *ACTA2* expression with the combination of SB431542 and OTR4120 compared to SB431542 alone indicates the presence of additional mechanism(s) of action of OTR4120.

Our bioinformatic analysis indicated that OTR4120 had a significant impact on gene expression in our WI-38 cell model. *SERPINE1* is a gene known to be associated with lung fibrosis [30, 31]: SERPINE1 inhibits tissue plasminogen activator (tPA) and urokinase (uPA), enzymes involved in the breakdown of fibrin clots [32]. By inhibiting these activators, SERPINE1 promotes the accumulation of fibrin and the development of fibrosis in the lung [33]. Additionally, *GREM1* and *GREM2* were also found to be downregulated after treatment with OTR4120. Both proteins inhibit BMP signalling, which is important for controlling cell growth and tissue repair. By inhibiting BMP signalling, GREM1 and GREM2 contribute to fibrosis development [34–36]. They also interact with TGF-β1 signalling, leading to enhanced fibroblast activation and increased production of extracellular matrix, which are key features of fibrotic tissue remodelling [34]. Ultimately, these actions result in the thickening and stiffening of lung tissue, characteristic of fibrosis. Upregulated genes by OTR4120 included *PTPRD* (*Protein Tyrosine Phosphatase Receptor Type D*) that plays a role in cell signalling pathways that regulate various cellular processes. In lung fibrosis, PTPRD may exhibit antifibrotic properties by dephosphorylating and inactivating signalling molecules involved in profibrotic pathways [37]. Another gene that was upregulated was *ALDH1A3* (*Aldehyde Dehydrogenase 1 Family Member A3*), which encodes for an enzyme responsible for metabolizing aldehydes, converting them into carboxylic acids [38]. In lung fibrosis ALDH1A3 acts as an antifibrotic agent by facilitating the detoxification of aldehydes, thereby preventing cellular damage and subsequent fibrotic responses [38]. However, the upregulation of certain profibrotic genes (AGT (Angiotensinogen) or C3 (Complement Component 3)) leads to the idea of a complex system, where it is the final balance of pro- and anti-fibrotic genes that represents the most important step for tissue repair [39–42]. Of note, RGTA® can stabilize growth factors, enhancing their half-life and thus impacting fibrosis resolution [43].

Recent findings suggest that during the alveolar repair process, Runx1+ cells expressing Cthrc1 are present and may play a beneficial role [44]. Interestingly, these cells share similarities with the pathological Cthrc1+ activated fibroblasts found associated with the lung fibrosis [45]. It is proposed that the abnormal persistence of these cells is associated with fibrosis. In this context, the observed effect of OTR4120 on the MYF signature, could potentially promote MYF differentiation towards a beneficial cell type. However, further studies are needed to define the transient state of the fibroblasts in both normal and pathological conditions.

Nevertheless, the overall effect of OTR4120 is positive. While excessive myofibroblast presence is typically linked to fibrosis, their controlled activation is crucial for effective tissue repair and wound healing. The positive outcomes across multiple models suggest that OTR4120 treatment induces a regulated myofibroblast response that can be translated into a healthy regeneration process. The observed positive effects support the idea that OTR4120 achieves a balance between promoting necessary fibrotic activity for repair and tissue regeneration while preventing excessive fibrosis that could lead to detrimental outcomes.

Furthermore, in the *in vivo* model of lung fibrosis induced by bleomycin administration in mice, OTR4120 demonstrated a remarkable ability to accelerate fibrosis resolution. It is important to note that, in this model, bleomycin was administered IP whereas other murine models where bleomycin is applied by another route might provide different results.

The results all together suggest that in human, OTR4120 could be effective not only in halting fibrosis progression like the approved drugs Nintedanib and Pirfenidone, but also in promoting the resolution of existing fibrotic lesions. In addition, the effects of OTR4120 to modulate tissue regeneration and reduce inflammatory cell infiltration further contribute to its therapeutic potential in fibrotic diseases.

Considering the modulation of the immune response, in the bulk RNA-Seq many immune-related genes were impacted by the OTR4120 treatment (*MMP1*, *TNFRSF12A* -*Tumor Necrosis Factor Receptor Superfamily Member 12A*-, *EGR1*-*Early Growth Response 1*). The negative enrichment of immune-related genes, including those involved in interleukin-4 and interleukin-13 signalling, suggests a suppression of immune responses in the OTR condition. This reduction in immune activity may contribute to a less inflammatory environment, potentially impacting processes such as tissue repair and fibrosis regulation, where immune pathways play a crucial role.

## Conclusion

These findings provide a solid foundation for further exploration and development of OTR4120 as a potential therapeutic approach for lung fibrosis. The broad effect of the OTR4120 that emerges from these data clearly pointed out the complexity of the mode of action, that need to be further explored in order to exploit all the potential of this promising drug. Over time, the healthy lung is prone to accumulating damage due to various injuries, which can lead to pathological fibrosis. We propose that OTR4120 has the capability to reverse the activation of myofibroblasts, which are primarily responsible for excessive extracellular matrix (ECM) deposition and increased tissue stiffness. By promoting the reversion of activated myofibroblasts to a healthier state, OTR4120 may facilitate tissue repair and regeneration, leading to the restoration of functional alveoli and overall lung function.

## Acknowledgement

Authors wish to thank Professor Emeritus Jean Francois Bernaudin (University Paris Nord) for his precious comments and advice.

## Authors contribution

MM, SC, DW, MMK, TP, EVP, AN, YD, NMC, DPG carried out the experiments. JZ, FC, AC, RP, MT, HW, CR and AG contributed to the collection of the human specimens, MM, YZ, XC, AL, SC, MMK; and SF contributed to the analysis of the results. MB, SR, LG, TP and JW contributed to the bio-informatic analyses. MM, DW, SR contributed to the imaging. MM, DB, CMC, SR, WS and SB designed the project, contributed to the analysis of the results and wrote the paper.

---

**Supplementary Figure 1.**
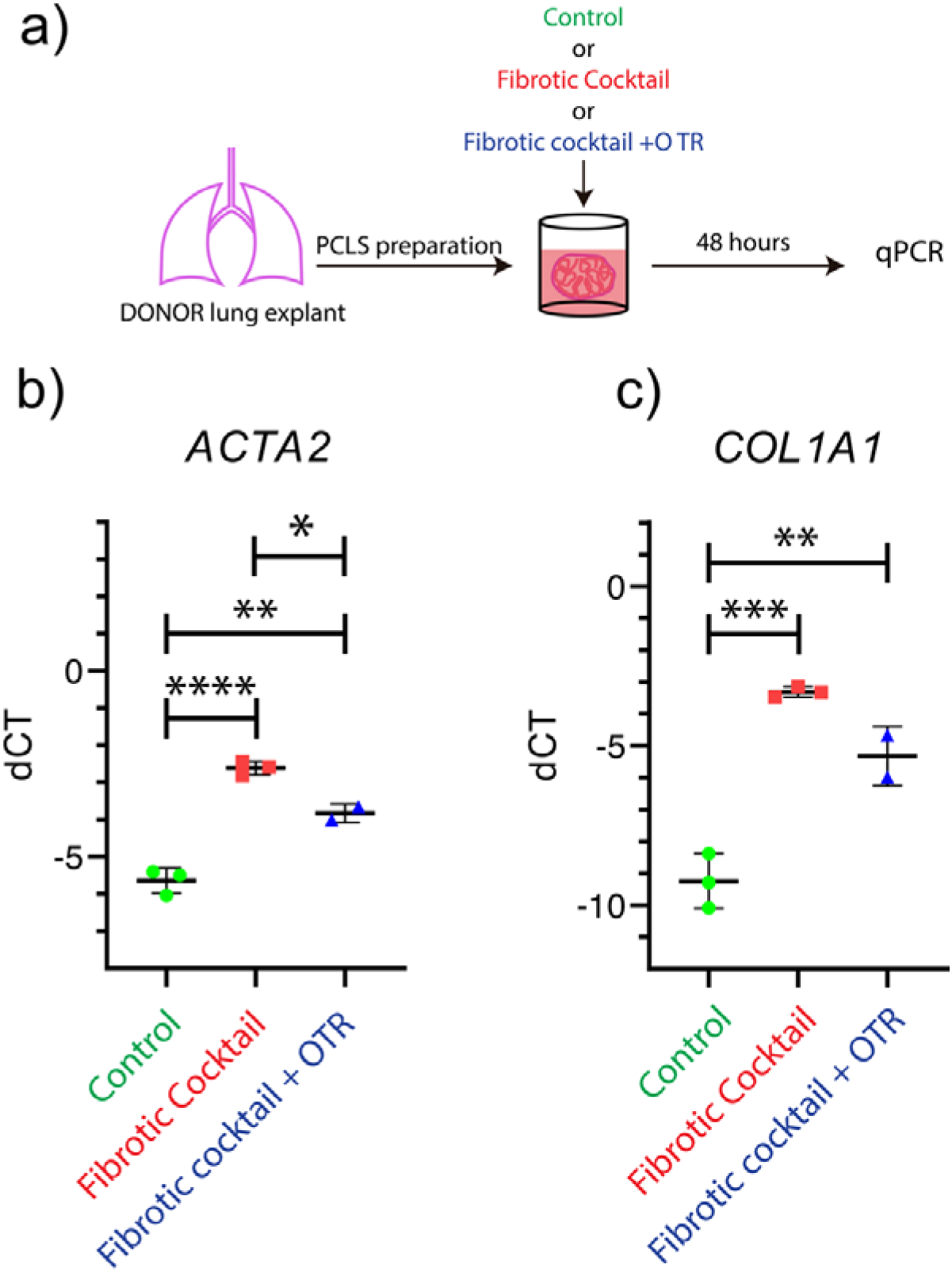
OTR4120 treatment of PCLS from human DONOR lungs grown in vitro in presence of a Fibrotic Cocktail leads to reduced expression of *ACTA2* and *COL1A1*. **(a)** Schematic representation of the experimental set up for the gene expression analysis in hPCLSs. **(b)** Gene expression analysis for MYF markers: *ACTA2* (left) and *COL1A1* (right).

**Supplementary Figure 2.**
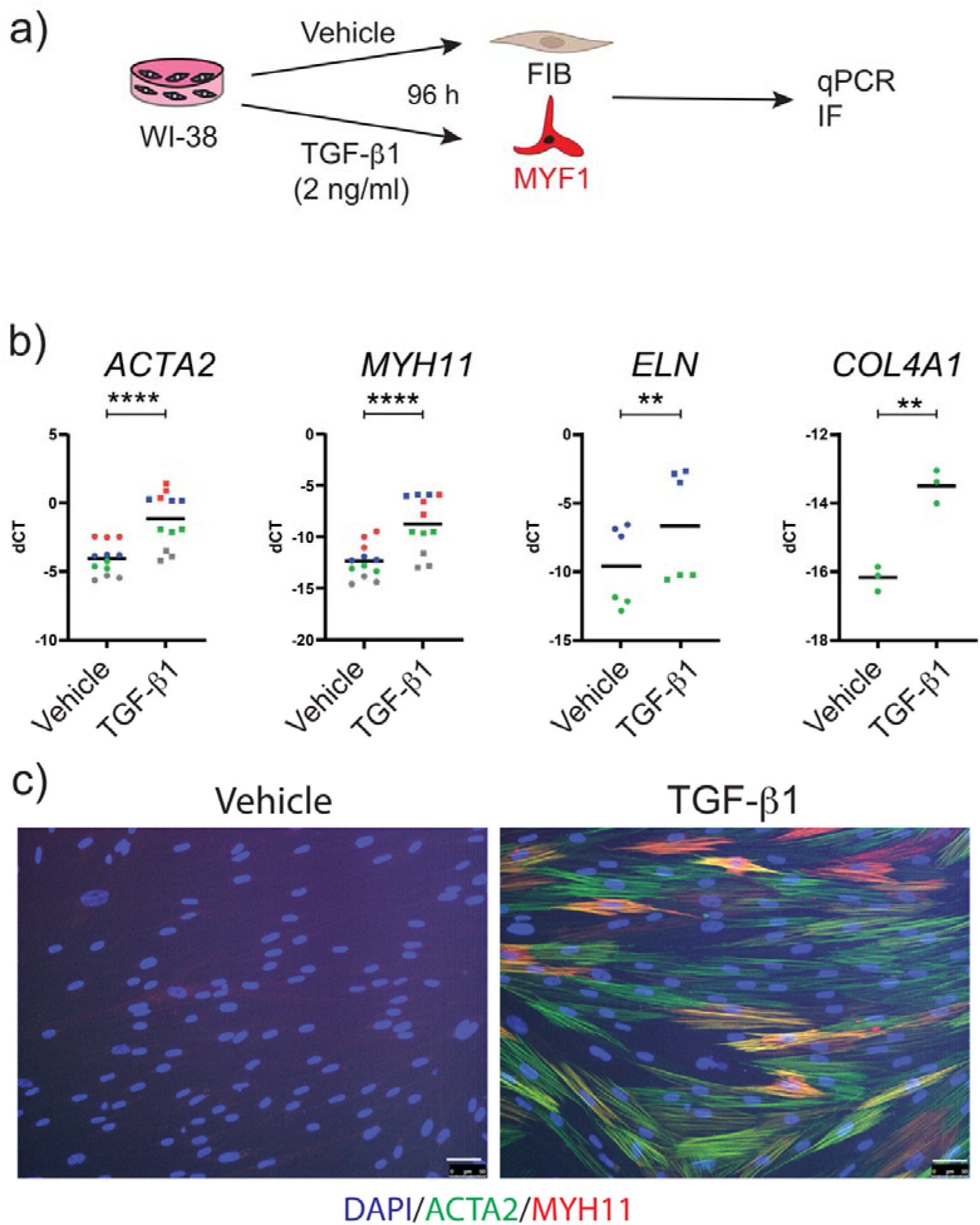
Impact of TGF-β1 treatment on WI-38 cells. **(a)** Schematic representation of the experimental set up. **(b)** Gene expression analysis for MYF markers in the vehicle control and TGF-β1 treated-WI-38: *ACTA2*, *MYH11*, *ELN*, *COL4A1*. **(c)** Immunofluorescence for the MYF markers ACTA2 (green) and MYH11 (red) in the vehicle control (left) and TGF-β1 treated-WI-38 (right). Nuclei are stained with DAPI. *P* values * *P* < 0.05; ** *P* < 0.01; *** *P* < 0.001; **** *P* < 0.0001. Scale bar c: 50 μm

